# ABCG transporter gene *PstABCG2* contributes to multidrug resistance in *Puccinia striiformis*

**DOI:** 10.1101/2025.08.20.671227

**Authors:** Fan Ji, Xinpei Gao, Yaning Liu, Ying Li, Lili Huang, Jun Guo, Gangming Zhan, Zhensheng Kang

**Affiliations:** State Key Laboratory for Crop Stress Resistance and High-Efficiency Production and College of Plant Protection, Northwest A & F University, Yangling 712100, P. R. China

## Abstract

The emergence of triadimefon resistance in *Puccinia striiformis* f. sp. *tritici* (*Pst*), the causal agent of wheat stripe rust, poses a growing threat to global food security. While the novel SDHI fungicide flubeneteram exhibits high efficacy against *Pst*, the molecular mechanisms governing resistance to both fungicides remain largely unknown. Here, we identify the ABC transporter PstABCG2 as a key mediator of multidrug resistance (MDR) through combined transcriptomic, genetic, and biochemical analyses. First, time-series transcriptomics revealed sustained upregulation of *PstABCG2* under fungicide stress, with silencing via in planta RNAi and HIGS significantly enhancing *Pst* sensitivity to both triadimefon and flubeneteram. Heterologous expression of *PstABCG2* in a *Fusarium graminearum* ΔFgABCG2 mutant restored fungicide tolerance, confirming its efflux function. Crucially, saturation mutagenesis and structural modeling pinpointed E1184 as a critical residue for fungicide binding: the E1184Y mutation disrupted triadimefon affinity (validated by MST/ITC, ΔG reduced from -7.1 to -6.5 kcal/mol) but minimally impacted flubeneteram binding due to adaptive halogen-bond networks. Furthermore, we discovered that the GATA-family transcription factor PstGATA directly activates *PstABCG2* expression by binding to promoter *cis*-elements, and silencing *PstGATA* phenocopied the hypersensitivity of *PstABCG2*-silenced isolates. The results may provide a theoretical foundation for elucidating multidrug resistance mechanisms in *Pst*, field resistance management, and novel fungicide target development.

## Introduction

Wheat, a staple crop vital to global food security, sustains 40% of the world’s population by supplying 20% of humanity’s dietary energy and protein requirements [1]. Across major wheat-producing areas worldwide, stripe rust disease, induced by the fungal pathogen *Puccinia striiformis* f. sp. *tritici* (*Pst*), frequently causes significant reductions in crop yield, posing a persistent challenge for agricultural communities internationally [2, 3]. Fungicide application serves as a key strategy in combating this disease, protecting yield potential and maintaining production consistency [4]. Nevertheless, the extensive and recurrent use of single-mode-of-action fungicides may accelerate the emergence of resistant *Pst* isolates, creating new control difficulties.

Triadimefon, the mainstay of stripe rust control, have been widely and exclusively used for over 50 years [5]. This sustained monocultural application has exerted strong selection pressure, driving resistance evolution and compromising chemical control efficacy. Our nationwide monitoring revealed 6.79% of *Pst* isolates displayed triadimefon resistance, with more insensitive isolates collected from *Pst* winter-increasing areas and northwest over-summering areas [6]. Monitoring of Chinese wheat rust populations revealed >5% triadimefon resistance in both *Puccinia triticina* (leaf rust) and *Puccinia graminis* (stem rust) [7, 8]. These findings highlight the emerging triadimefon resistance in rust pathogens, providing a foundation for investigating resistance mechanisms.

Triadimefon primarily target *Cyp51*, an essential cytochrome P450 enzyme that catalyzes lanosterol 14α-demethylation in the ergosterol biosynthesis pathway required for fungal cell membrane integrity [9, 10]. Currently, the primary mechanisms of triadimefon resistance in phytopathogenic fungi include: (1) single or multiple mutations in the *Cyp51* target gene, (2) *Cyp51* overexpression, (3) efflux-mediated detoxification by ABC (ATP-binding cassette) or MFS (major facilitator superfamily) transporters, and (4) compensatory overexpression of *Cyp51* homologs [11–13]. Our analysis of 486 Chinese *Pst* isolates identified the *Cyp51* Y134F mutation in moderately and some low-resistant isolates, but it was absent in other low-resistant isolates. *Cyp51* expression showed no resistance-associated changes, implying non-regulatory resistance mechanisms [6].

Flubeneteram is a novel succinate dehydrogenase inhibitor (SDHI) fungicide developed using the pharmacophore-linked fragment virtual screening (PFVS) method in China [14]. This fungicide offers advantages including high efficacy, low toxicity, and broad-spectrum activity. Previous studies have demonstrated that flubeneteram can control numerous crop diseases, such as rapeseed sclerotinia, soybean rust, melon powdery mildew, and corn rust [14]. Previous research from our group showed that flubeneteram exhibited excellent inhibitory activity against *Pst* [8]. However, the potential resistance mechanisms of *Pst* to this fungicide remain unclear.

Current studies demonstrate that ATP-binding cassette (ABC) transporters represent a large and functionally diverse family of membrane transport proteins in mammals, plants, and microorganisms. These transporters not only mediate toxic compound efflux but also play crucial roles in fungicide multidrug resistance (MDR) development in pathogenic fungi [15–17]. Research has established strong associations between MDR in various pathogens and specific ABC transporter functions. For instance, *Sclerotinia sclerotiorum* exhibits positive cross-resistance to dicarboximide fungicides (e.g., iprodione and procymidone), attributed to its ABC transporters’ capacity to efflux both phenylpyrrole and dicarboximide fungicides [18]. Furthermore, pathogens employ sophisticated transcriptional regulation to activate ABC transporters, thereby enhancing MDR. In *Botrytis cinerea*, for example, ABC transporters are positively regulated by transcription factor Mrr1 and contribute significantly to MDR [19]. However, the specific ABC transporter subtypes mediating fungicides resistance in *Pst* remain poorly characterized, and their transcriptional activation mechanisms in response to fungicide exposure require further investigation.

With the purpose of determining the biological roles of ABCG transporters in *Pst* and their fungicides and responsive transcriptional regulation, and uncovering key mechanisms of fungicide resistance mediated by these transporters, this study was conducted to achieve the following objectives: (i) identify ABCG transporter genes upregulated by fungicides through transcriptome analysis; (ii) employ host-induced gene silencing (HIGS) and RNA interference (RNAi) screening to identify functional ABCG transporters mediating fungicides resistance in *Pst*; (iii) characterize function through heterologous expression in *Fusarium graminearum* to demonstrate the fungicides efflux capability of candidate ABCG transporters; (iv) predict and analyze the conservation of key binding sites between core ABCG transporters and fungicides; and (v) screen for transcriptional regulators to identify specific factors controlling ABCG transporter expression, with subsequent mechanistic characterization of their activation/repression pathways. The results may provide a theoretical foundation for elucidating multidrug resistance mechanisms in *Pst*, field resistance management, and novel fungicide target development.

## Results

### Activation of ABC transporter pathways in *Pst* under triadimefon and flubeneteram treatment

To elucidate the molecular mechanisms underlying fungicide resistance, a time-series transcriptomic experiment was performed with *Pst*-infected wheat leaves subjected to triadimefon and flubeneteram treatment at 0, 12, 24, 48, and 120 h post-treatment, including three biological replicates per time point (totaling 15 samples per treatment group). Principal component analysis (PCA) of the transcriptomic data revealed that the 95% confidence ellipses of the triadimefon- and flubeneteram-treated samples overlapped substantially (Supplementary Fig 1A), indicating highly congruent transcriptomic responses to both fungicides. Based on this observation, both treatments were merged into a single experimental group (n=30) for DEG analysis against controls (n=15). Differential expression analysis identified 649 genes exhibiting significant changes (fold change >2, P < 0.05), of which 208 were upregulated and 441 downregulated (Supplementary Fig 1B). KEGG enrichment ranked the ABC transporter pathway among the top 20 enriched pathways (*P*=0.23; Figure 1A). Gene set enrichment analysis (GSEA) further confirmed significant upregulation of ABC transporters in the treated samples, with a normalized enrichment score (NES) of 1.52 (*P* < 0.05). Among the 14 pathways satisfying the gene set size criteria (15–500 genes), ABC transporters exhibited the most pronounced enrichment (Figure 1B), with sustained upregulation indicative of a synergistic stress response.

**Fig 1.**
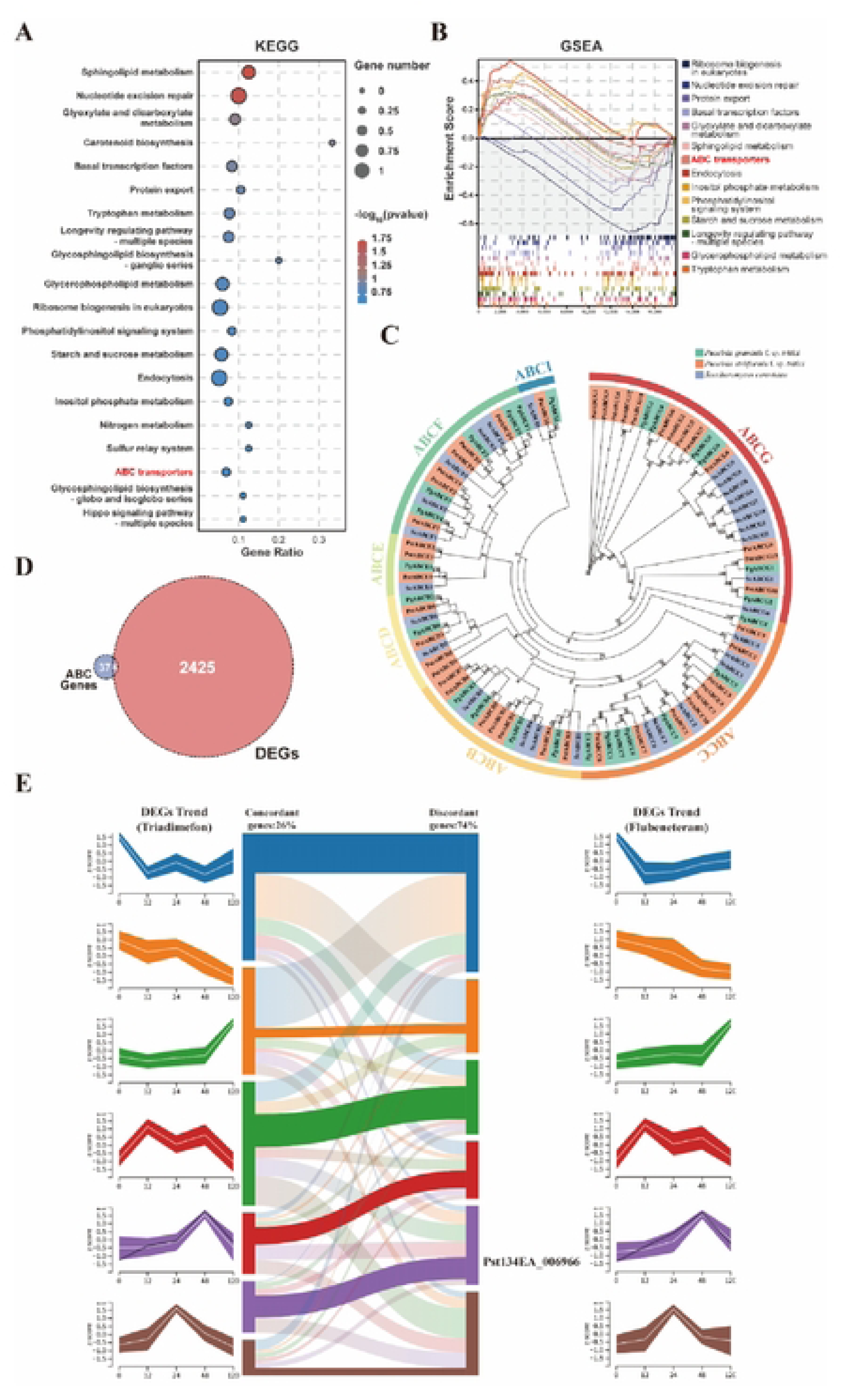
Regulation of ABC transporter pathways in *Pst* under triadimefon and flueneteram treatment. **(A)** KEGG enrichment of ABC transporters (top 20 pathways) in combined fungicide-treated vs. control groups. **(B)** GSEA plots: Top panel shows enrichment score (ES) curves; bottom depicts gene distribution. ABC transporters had the highest NES (1.52, *P* < 0.05). **(C)** Phylogenetic tree of 43 ABC transporters (ABC-G subfamily in red). **(D)** Venn diagram of six overlapping DEGs and ABC transporters. **(E)** OmicsTIDE time-series of *Pst134EA_006966* (*PstABCG2*) showing early upregulation (0–48 h).

Phylogenetic analysis identified 43 ABC transporters, including 13 (30.2%) from the ABC-G subfamily (Figure 1C). Venn diagram analysis revealed six overlapping genes between differentially expressed genes (DEGs) and the 43 bioinformatically identified ABC transporter genes (Figure 1D). Among these, four of which *Pst134EA_006966(PstABCG2)*, Pst134EA_007827*(PstABCG6)*, *Pst134EA_007831(PstABCG7)* and *Pst134EA_011091(PstABCG8)* belonged to the ABC-G subfamily, indicating these ABC transporter genes were also differentially expressed. This finding suggests these genes may directly mediate fungicide resistance. As shown in Supplementary Fig 2, compared to the control, the expression levels of the four ABC-G transporter genes were significantly upregulated at 24 and 48 h after the triadimefon and flubeneteram treatment. OmicsTIDE temporal analysis further revealed that the ABC transporter gene *Pst134EA_006966*(*PstABCG2*) was consistently upregulated within 0-48 hours under triadimefon and flueneteram treatments (Figure 1E), suggesting its potential role as a core responsive gene shared by both fungicides. This expression pattern aligns with the efflux function of ABC transporters, supporting its involvement in resistance development through active fungicide efflux. In conclusion, the ABC transporter gene *PstABCG2* was selected as the key target for subsequent in-depth research.

### Silencing *PstABCG2* significantly enhanced the sensitivity of *Pst* to triadimefon and flubeneteram

To investigate the role of *PstABCG2* in triadimefon and flueneteram sensitivity, we used a HIGS transient silencing approach to silence this gene. After 10 days of inoculation with PDS (lycopene dehydrogenase), obvious photobleaching symptoms were observed in wheat leaves, indicating that the target gene had been effectively silenced (Supplementary Fig. 3A). On this basis, *Pst* inoculation and fungicides treatment experiments were conducted on the silenced plants of the *PstABCG2*. The transcription levels of *PstABCG2* decreased by 46% - 67% in the silenced plants (Supplementary Fig. 3C-3D). Silencing *PstABCG2* significantly enhanced of the sensitivity of *Pst* in the triadimefon and flubeneteram treatment. Ten days after the two fungicides treatment, urediniospore production of the silenced isolates was substantially lower than in those in the controls (Supplementary Fig. 3B). To further confirm that silencing the three candidate genes can reduce the resistance of *Pst* to triadimefon, the mycelial development of silenced *PstABCG2* after fungicides treatment was observed under a microscope. Histological observations revealed that after 72 h of the fungicide treatment, mycelial growth was significantly inhibited in the silenced isolates (Supplementary Fig. 3D-3E).

To further conform the transient silencing results, RNAi was used to create a transgenic *Pst* isolate in the wheat variety Fielder. Following PCR testing and screening of positive plants in the T_0_ transgenic lines, two consecutive generations were propagated to produce T_3_ seeds. As shown in Fig. 2A, the triadimefon and flubeneteram sensitive levels of the T3 transgenic plants with RNAi silenced *PstABCG2* was significantly enhanced after inoculation with *Pst*. The expression levels of *PstABCG2* were reduced by 71% -78% in the RNAi lines (Fig. 2B). After 10 days of treatment with triadimefon, the biomass of *Pst* on RNAi transgenic lines was significantly lower than that in the control (Fig. 2C). For further determine whether the RNAi transgenic lines inoculated with *Pst* would affect the growth and development of hyphae after treatment with triadimefon and flubeneteram, the development of hyphae was observed and measured using a fluorescence microscope. As shown in Fig. 2D-2E, fluorescence microscopy revealed that hyphal growth was significantly inhibited in RNAi isolates after the fungicide treatment. These results confirmed that silencing *PstABCG2* through RNAi in Fielder wheat significantly enhanced *Pst* sensitivity to triadimefon and flubeneteram, consistent with the transient silencing results, which suggests that this gene play a key role in mediating sensitive to fungicides.

**Fig 2.**
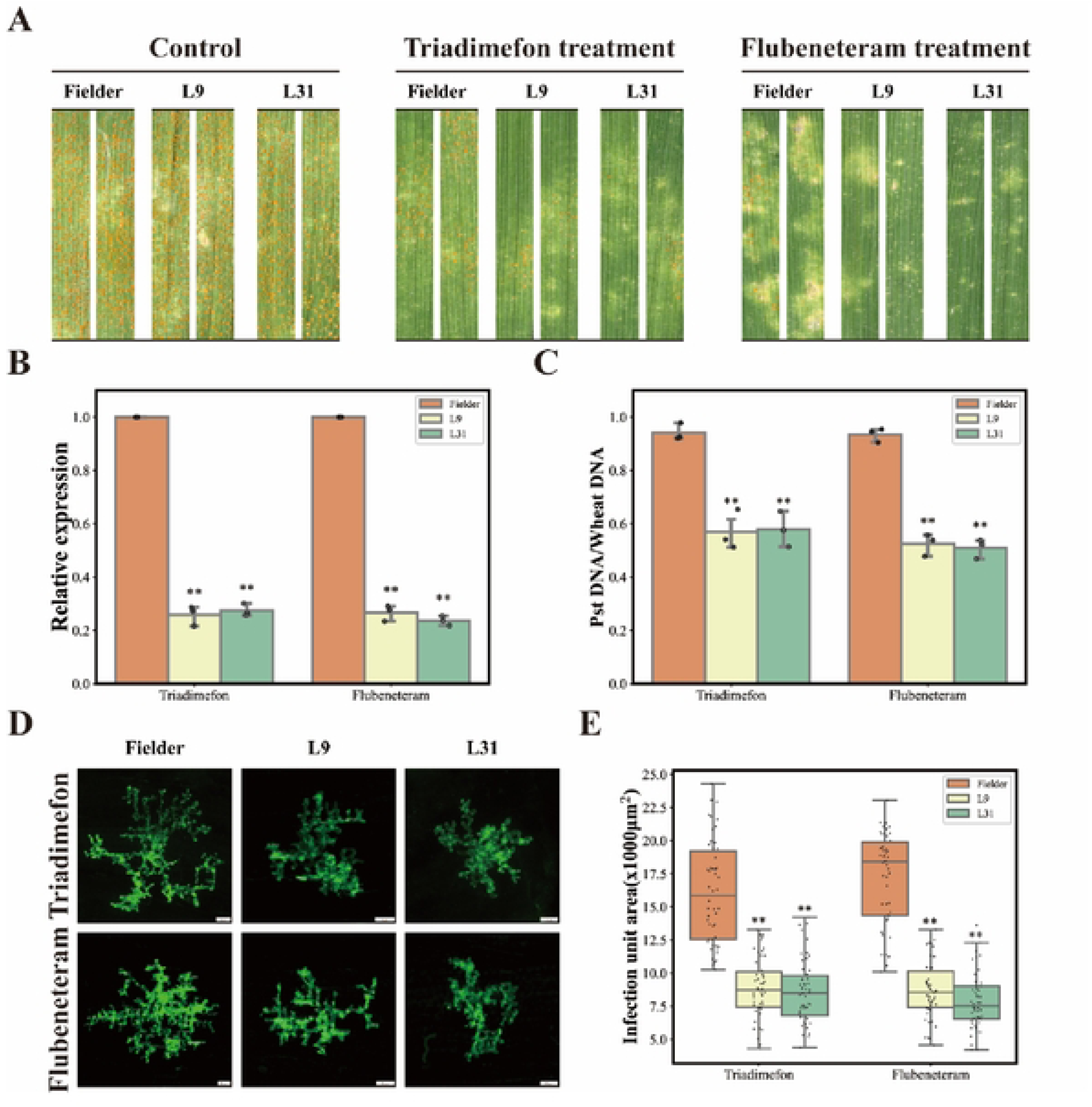
RNAi transgenic of *PstABCG2* plants enhance the sensitivity of *Pst* to triadimefon and flueneteram. **(A)** Disease phenotype of *Pst* inoculated with the RNAi plants under both fungicides and control treatments. **(B)** Silencing efficiency assay of *PstABCG2* in the RNAi plants under fungicides and control treatment. **(C)** Comparison of the biomasses of urediniospore produced on wheat leaves treated and non-treated with fungicides for 10 days. **(D)** Fluorescence microscopy observation of mycelial growth of *Pst* in wheat leaves after silencing of *PstABCG2* after 72 hours post inoculation (hpi) of fungicides treatment and non-treatment. **(E)** Comparison of the mycelial area of each infected unit in wheat leaves treated with fungicides and water at 72 hpi. Values are the mean ± standard deviation (SD) of three independent samples with 60 infection sites accounted. Single asterisk (*P* < 0.05) and double asterisks (*P* < 0.01) indicate a significant difference from the non-treated control according to the Student’s *t*-test.

### Heterologous overexpression of *PstABCG2* and it mutant in Δ FgABCG2 for triadimefon and flubeneteram sensitivity assay

To investigate the role of the *Fusarium graminearum* homologous gene *FGSG_04580* (*FgABCG2*) in fungicides sensitivity, we performed knockout and complementation of *FgABCG2* in the ΔFgABCG2 mutant, followed by triadimefon and flubeneteram sensitivity assays. As shown in Fig. 3A-3C, there was no significant difference in hyphal growth rate or conidia production between the *FgABCG2* knockout and complementation strains grown on PDA medium. However, the knockout mutant exhibited significantly higher sensitivity to triadimefon and flubeneteram compared to both the complementation and wild-type strains (Fig. 3D-3E). These findings suggest that the homologous gene *FgABCG2* mediates triadimefon and flubeneteram sensitivity in *Fusarium graminearum*.

**Fig 3.**
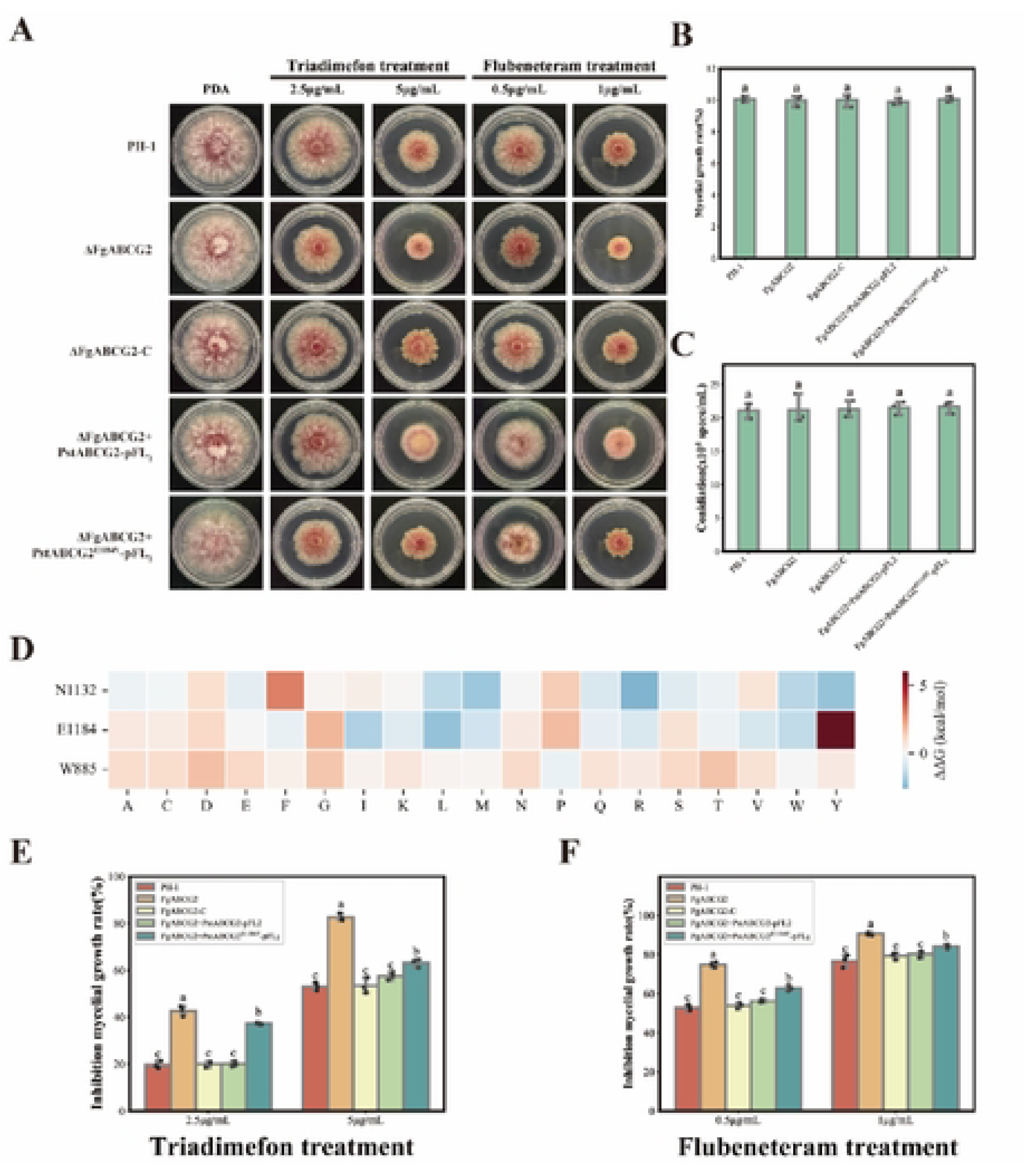
Sensitivity determination of PstABCG2 and its mutant heterologous overexpression in ΔFgABCG2 to triadimefon and flueneteram. **(A)** Phenotypic identification of PstABCG2 and its mutant heterologous overexpression in ΔFg ABCG2. Growth of *Fusarium graminearum* isolates on the normal potato dextrose agar medium (PDA), treatment medium (PDA + 2.5 and 5 μg/mL triadimefon) and treatment medium (PDA + 0.5 and 1 μg/mL flueneteram) for 3 days. **(B)** Comparison of mycelial growth rates of wild-type isolate PH-1, ΔFgABCG2, the complemented transformant ΔFgABCG2-C, PstABCG2 and its mutant heterologous overexpression in ΔFgABCG2 on potato dextrose agar (PDA) medium. Means and standard deviations (SD) were calculated from three repeats. **(C)** Comparison of conidial production of wild-type isolate PH-1, ΔFgABCG2, the complemented transformant ΔFgABCG2-C, PstABCG2 and its mutant heterologous overexpression in ΔFgABCG2 on PDA medium. Means and SD were calculated from three repeats. **(D)** Saturation mutagenesis analysis of key binding sites of PstABCG2 with triadimefon and flueneteram. The table is color-coded by correlation based on the color legend. Inhibition mycelial growth rate (%) of the wild-type isolate PH-1, ΔFgABCG2, the complemented transformant ΔFgABCG2-C, PstABCG2 and its mutant heterologous overexpression in ΔFgABCG2 to triadimefon **(E)** and flueneteram **(F)**. Bars with the same letter were not significantly different at *P*=0.05.

To assess the response of *PstABCG2* to triadimefon and flubeneteram, we heterologously overexpressed this gene in the ΔFgABCG2 mutant. The coding sequences of *PstABCG2* was cloned into the pFL2 vector and expressed in the ΔFgABCG2 mutant. The transformed strain was then cultured on PDA medium and PDA medium supplemented with 2.5 μg/mL and 5 μg/mL triadimefon, as well as PDA medium with 0.5 μg/mL and 1 μg/mL flubeneteram, to determine their sensitivity to triadimefon and flubeneteram under above conditions. As shown in Fig. 3A, overexpression of *PstABCG2* partially complemented the phenotype of the ΔFgABCG2 mutant. Specifically, under these two fungicides treatment, the overexpression strain restored growth defects in ΔFgABCG2.

Furthermore, saturation mutagenesis of the Vina-predicted binding sites in *FgABCG2* was conducted using FoldX to evaluate the effects of mutations on protein stability and fungicides binding affinity. As shown in Fig 3B, the substitution of amino acid residue 885 and 1132 with any other amino acid contributed minimally to protein structural changes. In contrast, the E1184Y mutation induced the most significant conformational alterations in the protein structure. We therefore generated complementation transformants by introducing site-directed mutations at nucleotides encoding amino acid residues 1184 of *PstABCG2* into the ΔFgABCG2 knockout mutant, and performed sensitivity assays on PDA and media supplemented with either triadimefon and flubeneteram. As shown in Fig 3D-3E, the overexpression strain of PstABCG2^E1184Y^ displayed lower phenotypic recovery against the ΔFgABCG2 mutant under triadimefon and flubeneteram stress compared to FgABCG2 mutant. Collectively, these results demonstrate that FgABCG2-the homologous gene of *Puccinia striiformis* PstABCG2 in *Fusarium graminearum*-similarly mediates sensitivity to both tested fungicides. Furthermore, heterologous overexpression of *PstABCG2* and its mutant PstABCG2^E1184Y^ in *Fg* revealed that transformants expressing PstABCG2^E1184Y^ exhibited heightened sensitivity to both fungicides compared to those expressing wild-type PstABCG2. This indicates that the E1184Y mutation plays a crucial role in the fungicide-binding process of this transporter protein.

### Determination of affinity for triadimefon and flubeneteram by purified proteins of PstABCG2 and PstABCG2^E1184Y^

To assess the affinity and transport capability of the PstABCG2 and it mutant for triadimefon and flubeneteram, molecular docking analysis was performed. Molecular docking analysis revealed distinct binding modes between flubeneteram (Flu) and triadimefon (Tri) with the PstABCG2 transporter. As shown in Fig. 4A, Tri exhibited moderate binding affinity (ΔG = -7.1 kcal/mol) through a specific hydrogen-bond network involving its triazole ring forming a hydrogen bond with W885 (3.0 Å), dual hydrogen bonds with N1132 (3.1 Å and 3.7 Å), and π-π stacking with F1191 (3.8 Å). In contrast, Flu demonstrated stronger binding (ΔG = -9.1 kcal/mol) via a multifaceted interaction network, including a halogen-bond triangle formed by fluorine atoms with E1184 (3.0 Å), E884 (3.1 Å) and D611 (2.8 Å), carbon-hydrogen bonds with W885 (2.9 Å) and N1132 (3.2 Å), and π-π stacking with F1191 (4.5-4.6 Å) (Fig. 4B). The E1184Y mutation caused significant protein destabilization (ΔΔG = +5.98 kcal/mol) but differentially affected fungicide binding: Tri lost critical interactions due to W885 conformational changes despite gaining a π-anion interaction with D611 (4.3 Å), resulting in reduced binding affinity (ΔG = -6.5 kcal/mol) (Fig. 4C), while Flu maintained high affinity (ΔG = -8.8 kcal/mol) through binding site adaptations where Y1184 preserved halogen bonding (3.0 Å), W885 interactions shifted from carbon-hydrogen to halogen bonds (both 2.9 Å), and a new hydrophobic contact with D611 (3.8 Å) emerged(Fig. 4D). These findings demonstrate how specific residue mutations can selectively impact fungicide binding through distinct molecular interaction mechanisms.

**Fig 4.**
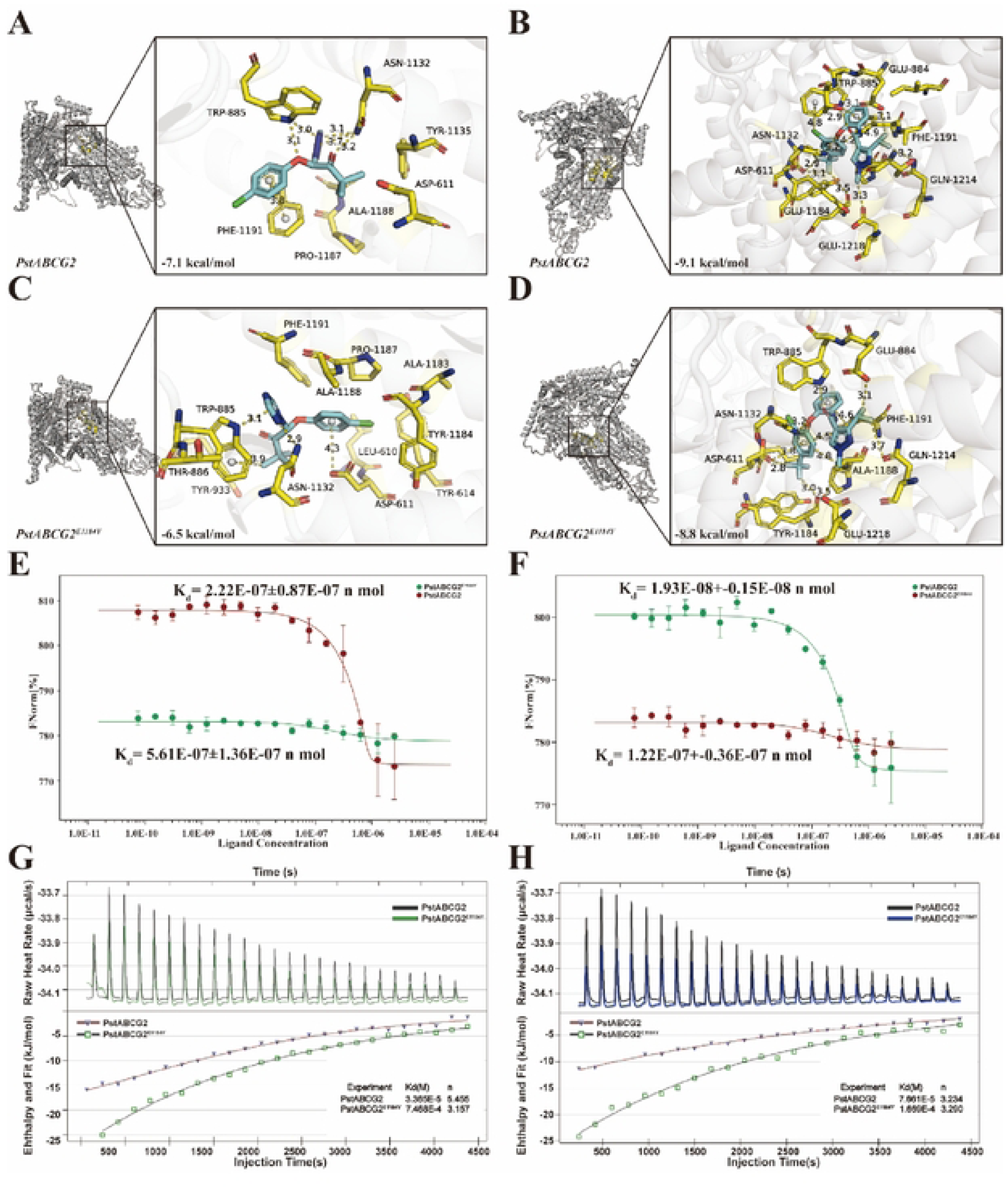
Molecular docking, microscale thermophoresis (MST) and isothermal titration calorimetry (ITC) analyses for binding interactions of PstABCG2 and PstABCG2^E1184Y^ with triadimefon and flueneteram. Molecular docking of PstABCG2 with triadimefon **(A)** and flueneteram **(B)**. Molecular docking of PstABCG2 ^E1184Y^ with triadimefon **(C)** and flueneteram **(D)**. Comparison of triadimefon **(E)** and flueneteram **(F)** binding affinity between PstABCG2 and PstABCG2^E1184Y^ mutant using MST. The binding curves fit the data points to the standard K_d_-fit function. Mean and standard deviations were calculated with the data from three independent replicates. Comparison of triadimefon **(G)** and flueneteram **(H)** binding affinity between PstABCG2 and PstABCG2^E1184Y^ mutant using ITC. K_d_, the dissociation constant. n, stoichiometric ratio.

To further validate these findings, we used Microscale Thermophoresis (MST) and Isothermal Titration Calorimetry (ITC) to measure the binding affinity of these two proteins for triadimefon and flubeneteram. As shown in Supplementary Fig. 4, SDS-PAGE confirmed that purified proteins with the target band sizes were obtained for further affinity analysis. MST results revealed that both PstABCG2 and PstABCG2^E1184Y^ bound triadimefon, with dissociation constants (Kd) of 2.22 ×10^-7^ nM and 5.61×10^-7^ nM, respectively (Fig. 4E). Similarly, PstABCG2^E1184Y^ showed a significantly lower dissociation constant for flubeneteram than PstABCG2, with Kd values of 1.22 ×10^-7^ nM and 1.93 ×10^-8^ nM, respectively (Fig. 4F). Additionally, ITC measurements confirmed these findings, showing that PstABCG2 and PstABCG2^E1184Y^ are capable of binding triadimefon and flubeneteram, with varying dissociation constants. The binding affinity of PstABCG2^E1184Y^ was significantly lower than their wild-type counterparts. ITC also provided the stoichiometric ratio (n), which reflects the number of fungicides bound per protein molecule. As shown in Fig. 4G, PstABCG2 exhibited a significantly higher triadimefon binding affinity (Kd = 3.37×10^-5^ nM, n = 5.46) and stoichiometric ratio compared to PstABCG2^E1184Y^ (Kd = 7.47×10^-4^ μM, n = 3.16). Similarly, PstABCG2 showed a significantly higher affinity for flubeneteram (Kd = 7.66×10^-5^ nM, n = 4.23) than PstABCG2^E1184Y^ (Kd = 1.67×10^-4^ nM, n = 3.29) (Fig. 4H). This suggests that base mutations in PstABCG2 impair fungicide transport capacity, which may contribute to the increased sensitivity to triadimefon and flubeneteram.

### PstGATA was responsible for activating *PstABCG2* via binding on its promoter in *Pst* in response to fungicides treatment

Given that PstABCG2 is involved in the efflux of fungicides in *Pst*, we further identified hub regulators regulating PstABCG2 expression under triadimefon and flubeneteram treatment using WGCNA (Fig. 5B). The expression data of all genes encoding transcription factors and representative ABC transporters were collected for coexpression regulation network construction based on Pearson algorithm. Typically, the soft threshold was set to 5 (R^2^ = 0.85) for constructing a scale-free network, and 5 gene modules were categorized by hierarchical clustering. Among 5 modules, MEturquoise exerted the highest correlation value with PstABCG2, thus the transcription factors from this module were identified as candidate regulators for PstABCG2 expression (Fig. 5B). Then, all genes from MEturquoise were filtered using P < 0.05 and Pearson coefficient > 0.85 to construct the topological network. Totally, 48 genes encoding transcription factors from MEturquoise module were identified to involve in regulating PstABCG2 expression in *Pst* in response to triadimefon and flubeneteram treatments. We noted that most of these genes encoding transcription factors were highly expressed in *Pst* following fungicides treatment at 24 h, demonstrating their importance in regulating defense response of *Pst* to triadimefon and flubeneteram (Fig. 5A). Notably, *Pst134EA_030708* (*PstYAP1*), *Pst134EA_007617* (*PstGATA*), *Pst134EA_015014* (*PstC2H2-1*), *Pst134EA_009356* (*PstZFP36*) and *Pst134EA_025399* (*PstC2H2-2*) of all genes from MEturquoise exerted the highest positive relationship (Pearson coefficient = 0.75∼0.95) with PstABCG2 in the topological regulation network, thus identified as hub transcription factors regulating PstABCG2 expression under fungicides treatment (Fig. 5B-5C). Subsequent analysis using Jasper database predicted 3 cis-elements for GATA within PstABCG2 promoter, indicating the regulatory function of PstGATA for PstABCG2 expression (Fig. 5D).Then, yeast one-hybrid (Y1H) and LUC report assays demonstrated that PstC2H2-1 directly bound to the PstABCG2 promoter, thereby activating its expression in vivo (Fig. 5E-5F). Eventually, EMSA confirmed the binding activity of PstGATA to the P2 site containing a GATA-box motif (Fig. 5G). Thus, these results suggested that the higher level of PstGATA activated PstABCG2 expression via binding on its promoter under triadimefon and flubeneteram treatment.

**Fig 5.**
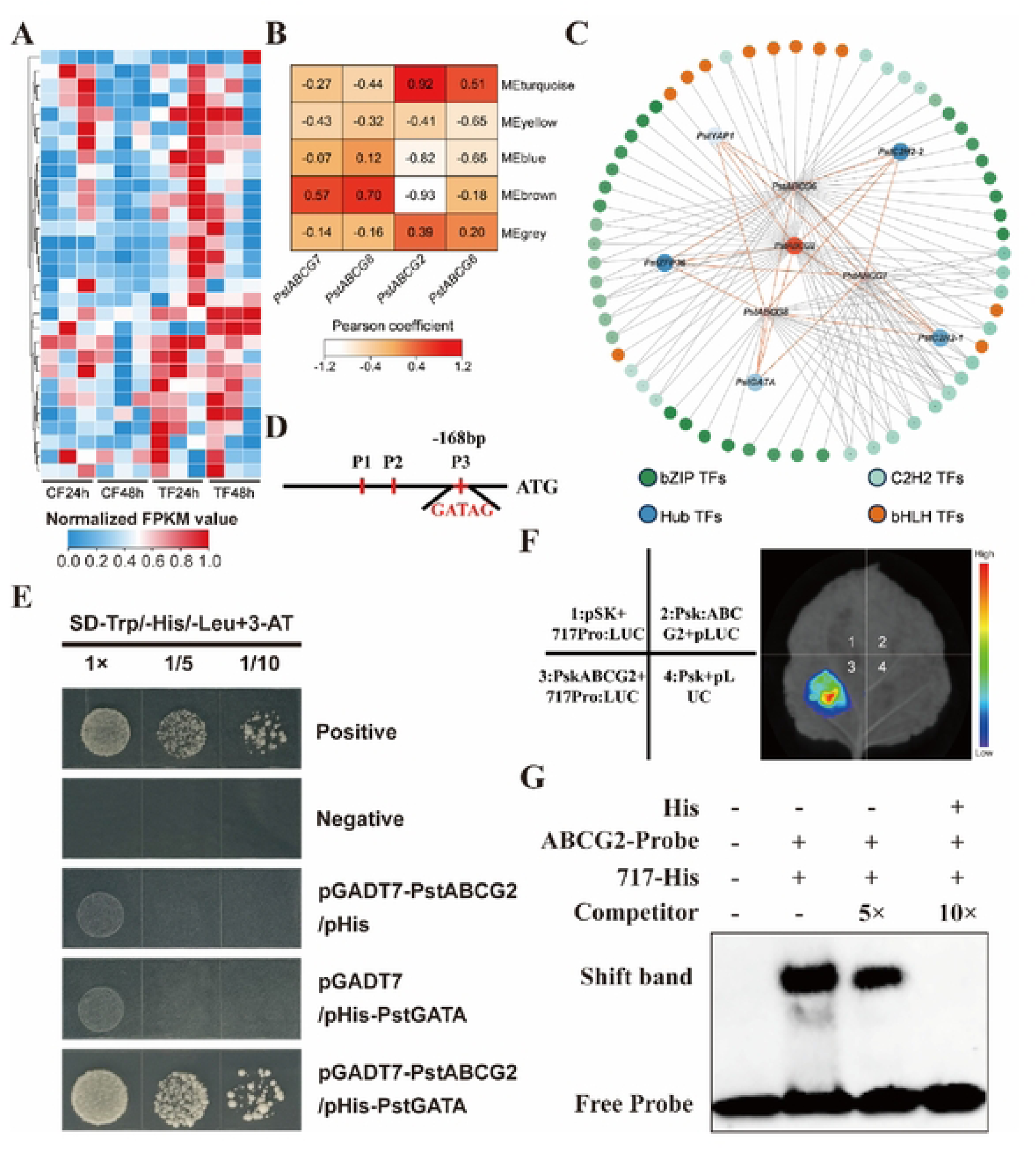
PstGATA was responsible for activating *PstABCG2* via binding on its promoter in *Pst* in response to fungicides treatment. **(A)** Heat maps of the relative expression abundances of genes encoding transcription factors from the regulation network for mediating ABC expressions in *Pst* following fungicide treatment. The scale bar showed the mean values of normalized FPKM in each experimental replicate. **(B)** Heatmap displaying the Module-trait relationships. Each row represents a module eigengene, and column represents a trait. Each cell contains the corresponding correlation and p value. The table is color-coded by correlation based on the color legend. **(C)** The weight network of the significant transcription factors involved in regulating genes encoding ABC transporter of *Pst* in response to fungicide treatment. The plot color represented the type of transcription factors in the network. **(D)** Identification of the *cis*-elements in *PstABCG2* promoter using Jasper database. **(E)** Yeast one-hybrid assay detects the binding affinity of PstGATA on *PstABCG2* promoter. **(F)** LUC fluorescence reporter assay displays the binding activities of PstGATA on *PstABCG2* promoter in *vivo*. **(G)** EMSA assay examines ability of PstGATA binding to the GATA-box (GATAG) of *PstABCG2* promoter. Biotin-labeled probes were incubated with PstGATA-His proteins, and the free and bound DNAs were separated on an acrylamide gel. As indicated, unlabeled probe was used as competitor.

### PstGATA transcriptionally activates PstABCG2 to reduce the sensitivity of *Pst* to fungicides

To investigate the role of PstGATA in triadimefon and flueneteram sensitivity, we used a HIGS transient silencing approach to silence this gene. After 10 days of inoculation with PDS (lycopene dehydrogenase), obvious photobleaching symptoms were observed in wheat leaves, indicating that the target gene had been effectively silenced (Fig. 6A). On this basis, *Pst* inoculation and fungicides treatment experiments were conducted on the silenced plants of the *PstGATA*. The transcription levels of *PstGATA* decreased by 42% - 62% in the silenced plants (Fig. 6B). Silencing *PstGATA* significantly enhanced of the sensitivity of *Pst* in the triadimefon and flubeneteram treatment. Ten days after the two fungicides treatment, urediniospore production of the silenced isolates was substantially lower than in those in the controls (Fig. 6C). To further confirm that silencing the three candidate genes can reduce the resistance of *Pst* to triadimefon, the mycelial development of silenced *PstGATA* after fungicides treatment was observed under a microscope. Histological observations revealed that after 72 h of the fungicide treatment, mycelial growth was significantly inhibited in the silenced isolates (Fig. 6D-6E). The findings indicate that PstGATA transcriptionally upregulates PstABCG2, consequently decreasing *Pst*’s sensitivity to fungicides.

**Fig 6.**
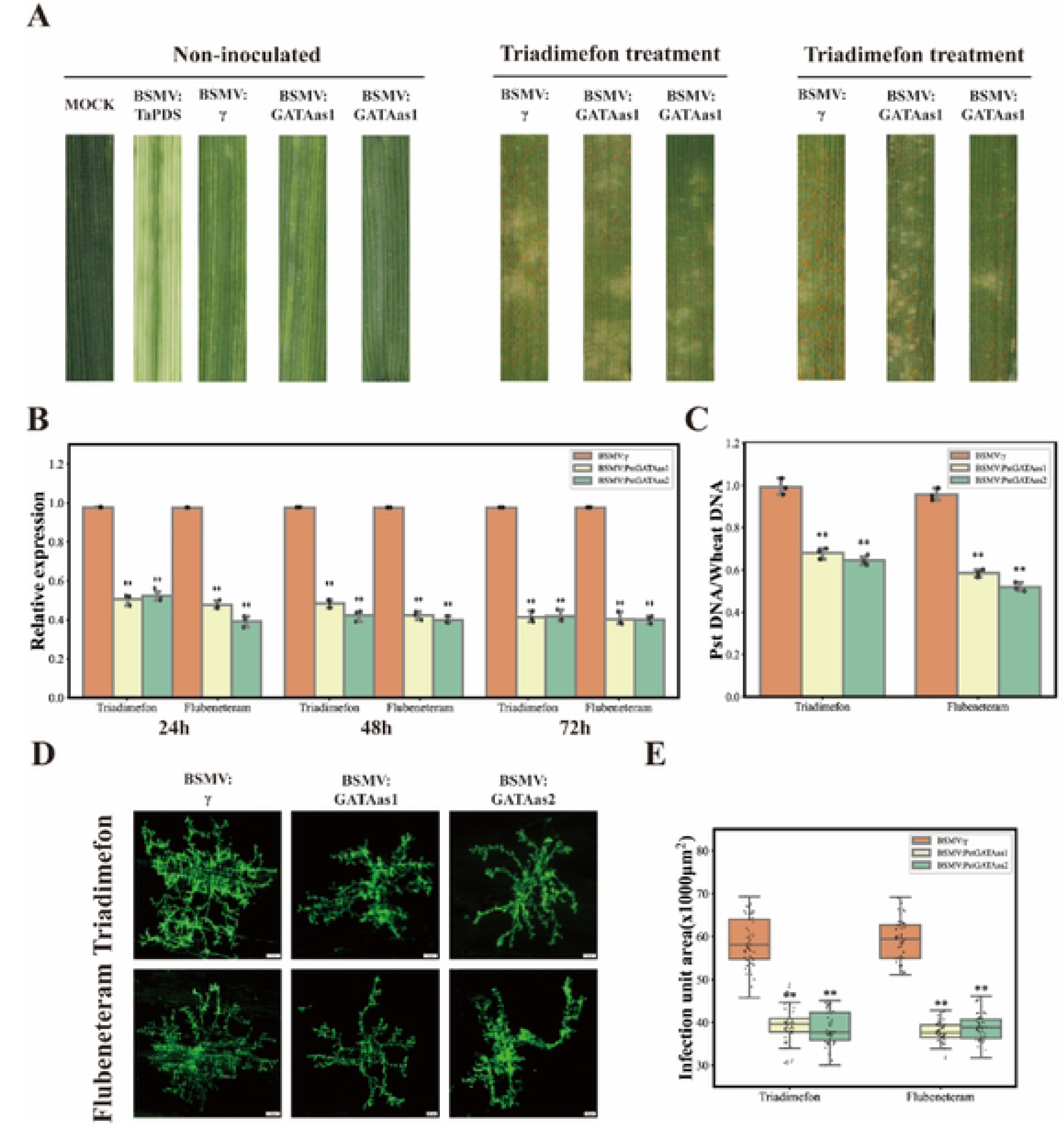
Silence of *PstGATA* enhances the sensitivity of *Pst* to triadimefon and flueneteram by BSMV-HIGS. **(A)** Disease phenotype of *Pst* inoculated with the fourth leaves of Sumon 11 that fully displayed viral symptoms under both triadimefon and flueneteram treatments. **(B)** Silencing efficiency assay of *PstGATA* in the HIGS plants under triadimefon and flueneteram treatments. RNA samples were isolated from the fourth leaves of knock-down and control plants at 24, 48 and 72 hpi of triadimefon and flueneteram treatments. The *PstEF* gene was used as an internal control. **(C)** Urediniospore biomass on wheat leaves treated with triadimefon and flueneteram at 10 days post-treatment. **(D)** Fluorescence microscopy observation of mycelial growth of *Pst* in wheat leaves after silencing of *PstGATA* after 72 hpi of triadimefon and flueneteram treatment. **(E)** Comparison of the mycelial area of each infected unit in wheat leaves treated with triadimefon and flueneteram at 72 hpi. Values represent the mean ± standard deviation of three independent samples with 60 infection sites. Single asterisk (*P* < 0.05) and double asterisks (*P* < 0.01) indicate a significant difference from the water control based on the Student’s *t*-test.

## Discussion

Wheat stripe rust, caused by *Puccinia striiformis* f. sp. *tritici*, represents a major Category I crop disease in China and poses a significant threat to global wheat production [4]. Triazole fungicides such as triadimefon have been primary control agents for over five decades. However, prolonged exclusive use has led to the emergence of resistance, as evidenced by our nationwide monitoring, which detected resistant isolates in 6.79% of surveyed populations, predominantly in key wheat-growing regions of China [6]. This emergence not only reduces the efficacy of disease control but also increases both economic and environmental burdens. Flubeneteram, a novel and effective fungicide candidate for controlling wheat stripe rust, has not yet been registered for this plant disease management [8]. Unraveling the molecular basis of fungicides resistance in *Pst* is therefore critical for developing sustainable management strategies and guiding next-generation fungicide discovery to combat this evolving threat.

The limited availability of viable drug targets in fungal pathogens makes control of fungal diseases in humans, animals, and plants particularly challenging [20]. This challenge is compounded by the rise of drug resistance. In plant pathogens, ABC transporters are among the most well-characterized mediators of fungicide resistance. These transporters exhibit broad substrate specificity across mammals, plants, and microorganisms [21], and play central roles in efflux processes and multidrug resistance (MDR) development. A systematic understanding of ABC transporter structure, function, and regulation is therefore essential to elucidate MDR mechanisms in plant pathogens and to design effective fungicide resistance management strategies.

Active efflux of fungicides constitutes a primary resistance mechanism, reducing intracellular concentrations of toxic compounds [22]. Genomic analyses of resistant fungal strains indicate that overexpression of efflux pump genes is a major contributor to antifungal resistance [23]. Laboratory studies demonstrate that fungicide exposure can directly induce efflux pump upregulation, confirming their essential role in fungal adaptation to fungicide. The primary fungicides efflux pumps fall into two major classes: ATP-binding cassette (ABC) transporters and Major Facilitator Superfamily (MFS) proteins [24]. In plant pathogenic fungi, ABC transporters are central to multidrug resistance [25, 26]. Notably, recent studies on *Saccharomyces cerevisiae* have elucidated MDR ABC transporter networks, providing valuable insights into their regulatory mechanisms [27].

Similar studies in *Rhizoctonia solani* demonstrate the functional importance of MDR ABC transporters in phytopathogens [23]. Research on pleiotropic drug resistance (PDR) in *Pst* has revealed critical knowledge gaps regarding its ABC transporters and regulatory networks. No definitive PDR-type ABC transporters have been identified in this wheat stripe rust pathogen, and their functional mechanisms remain unknown. Addressing this gap through systematic investigation of *Pst* ABC transporters is essential for: (1) deciphering fungal resistance evolution, (2) guiding novel fungicide development, and (3) optimizing resistance management strategies. These advances are urgently needed to combat the escalating threat of fungicide resistance in wheat rust pathogens. This study identified a total of 43 ABC transporter proteins through phylogenetic analysis, among which 13 (30.2%) belong to the ABC-G subfamily, a proportion significantly higher than that of other subfamilies (e.g., ABC-A/B/C). The ABC-G subfamily (also known as the PDR subfamily) plays a central role in fungal drug resistance, which is highly consistent with previous findings. In various pathogenic fungi, ABC-G transporters are known to directly mediate resistance to fungicides by efflux mechanisms, suggesting that *Pst* may employ a similar strategy in response to triadimefon and flueneteram stress. Notably, ABC-G members are typically localized to the plasma membrane and are responsible for the efflux of hydrophobic compounds. Both triadimefon (logP = 2.8, a triazole) and flueneteram (logP = 4.8, an SDHI)-exhibit hydrophobic characteristics, implying that ABC-G proteins may directly contribute to resistance formation by effluxing these compounds.

Through integrated molecular docking, mutational scanning, and experimental validation, this study systematically deciphered the differential effects of the E1184Y mutation on PstABCG2 binding to triadimefon and flueneteram. The stringent geometric constraints of triadimefon’s binding mode made it particularly vulnerable to mutation-induced disruption, as evidenced by a substantial reduction in binding enthalpy (ΔH: -8.2 → -5.7) measured by ITC. This energetic penalty correlates with the breakdown of critical hydrogen bonds, compounded by the marginal contribution of a weak π-anion interaction with D611 (4.3 Å) that appears structurally observable but thermodynamically insignificant under physiological conditions. In striking contrast, flueneteram demonstrated remarkable resilience to the E1184Y mutation through a sophisticated multi-tiered adaptation mechanism: (1) The core halogen-bonding network (E1184/Y1184-E884-D611) maintained optimal geometry (3.0±0.2 Å) via remarkable conformational plasticity; (2) Dynamic sidechain rearrangements of W885 and N1132 enabled microenvironmental adaptation; and (3) A distinctive entropy-driven compensation (positive ΔS) emerged through mutation-induced conformational selection. This elegant “conformational selection-induced fit” paradigm not only explains flueneteram’s mutational resistance but also establishes a novel framework for designing next-generation fungicides with enhanced durability against resistance mutations. The comparative analysis reveals fundamental principles in protein-ligand adaptation - where triadimefon’s rigid binding suffers catastrophic failure upon mutation, flueneteram’s dynamic interaction network employs coordinated structural adjustments to maintain binding competence, providing a blueprint for engineering mutation-resistant agrochemicals. However, the regulatory networks governing ABC transporters exhibit remarkable complexity, involving multiple layers of control.

The transcriptional network governing multidrug resistance (MDR) in fungal pathogens is highly complex and involves multiple regulators, many of which remain unidentified. In the model yeast *Saccharomyces cerevisiae*, the well-characterized zinc finger transcription factors Pdr1 and its paralog Pdr3 orchestrate the pleiotropic drug response [28, 29]. They function as both activators and repressors by binding to pleiotropic drug response elements (PDREs) within the promoter regions of ABC transporter genes, such as *PDR5*, *PDR10*, and *PDR15* [29]. Studies in pathogenic fungi reveal diverse transcriptional regulators of MDR. In *Aspergillus fumigatus*, loss of the transcription factor ZfpA induces expression of the ABC transporter gene *atrF*, thereby altering azole susceptibility [30]. Similarly, in *Botrytis cinerea*, the transcription factor Mrr1 positively regulates *BcatrB* expression; notably, the V575M point mutation in Mrr1 leads to constitutive upregulation of *BcatrB* in the MDR1 strain [31–33]. Conversely, an unexpected negative regulatory role was identified for the GATA-type transcription factor Gln3 in *Candida albicans*. Gln3 suppresses the expression of the ABC transporter genes *CDR1* and *CDR2*, as well as their transcriptional regulator *PDR1*. Consistent with this, deletion of *GLN3* decreases fluconazole susceptibility [34].This study provides the first evidence that a GATA-family transcription factor (PstGATA) regulates fungicide sensitivity in *Pst* by positively regulating the ABC transporter gene *PstABCG2*. Identifying novel MDR regulators or mechanisms involving xenobiotic efflux pathways in plant pathogens may facilitate the development of new antifungal strategies.

In summary, this study first identified *PstABCG2* as a candidate ABC transporter implicated in multidrug resistance through transcriptomic analysis and experimental validation in *Pst*. Functional characterization via RNAi and heterologous overexpression confirmed its role in mediating sensitivity to triadimefon and flueneteram. Subsequently, AlphaFold3-guided saturation mutagenesis pinpointed critical fungicide-binding residues (E1184) within PstABCG2. *In vitro* biophysical validation by microscale thermophoresis (MST) and isothermal titration calorimetry (ITC) demonstrated significantly attenuated binding affinity in mutated variants, directly linking these residues to fungicide interaction. Finally, yeast one-hybrid (Y1H) screening coupled with electrophoretic mobility shift assays (EMSA) revealed that the transcription factor PstGATA positively regulates *PstABCG2* expression by binding specific *cis*-elements in its promoter region. Our results may contribute to the understanding of the mechanism of multidrug resistance in *Pst*, and also provide new approaches for the chemical control of plant disease.

## Materials and Methods

### Fungal isolate, wheat cultivar, growth conditions and fungicide

Three wheat cultivars-Mingxian169 (MX169), Suwon 11, and Fielder-were employed in this study. Seedlings were grown, inoculated with *Pst* urediniospores, and maintained post-inoculation under consistent conditions following established protocols [35]. Triadimefon (97.5% active ingredient) was procured from Xi’an Hytech Agrochemicals Co., Ltd. (Shaanxi, China). Flubeneteram (98% active ingredient) was provided by Dongguan Pesticide Research Co., Ltd (Dongguan, Guangdong, China).

### Cloning and Sequence analysis

The *PstABCG2* gene was PCR-amplified from *Pst* genomic DNA (strain 134E36_v1_pri; locus ID: Pst134E_006966). Genomic DNA was extracted from urediniospores using the CTAB method [36], and total RNA was purified from infected wheat leaves using the Quick RNA Isolation Kit (Huayueyang Biotechnology, China). Protein domains were predicted using HMMER (EMBL-EBI), and phylogenetic analysis was performed in MEGA 7.0 using the neighbor-joining method. Viral protein sequences used for phylogenetic reconstruction are listed in Supplemental Table 1.

### Plasmid construction

Recombinant BSMV-HIGS vectors were generated following Liu et al. [37]. Target-specific *PstABCG2* cDNA fragments for host-induced gene silencing (HIGS) were predicted using SiRNA Finder (Si-Fi) and cloned into the BSMV-γ vector via NotI and PacI restriction sites.

For RNAi-mediated gene silencing in wheat, *PstABCG2* cDNA fragments were inserted into the Gateway-compatible plasmid pC336 (Ubi:GWRNAi:Nos) [38].

For overexpression assays, the *PstABCG2* coding sequence was PCR-amplified and ligated into the *XhoI*-linearized pFL2 vector containing a geneticin resistance marker and GFP tag [39].

For protein purification, *PstABCG2* and its mutant variant (*PstABCG2^E1184Y^*) were cloned into *EcoRI/XhoI* sites of the pET28a vector, producing N-terminal His-tagged fusion proteins ( PstABCG2-His and PstABCG2 ^E1184Y^ -His).

For yeast one-hybrid (Y1H) assays, the *PstABCG2* promoter sequence was PCR-amplified with primers containing *EcoRI/XhoI* sites, purified, and cloned into pAbAi (Clontech) upstream of the Aureobasidin A resistance gene (AbAR). The PstGATA transcription factor cDNA was cloned into pGADT7-AD (Clontech) containing the GAL4 activation domain. All constructs were verified by Sanger sequencing.

For firefly luciferase reporter assay, *PstABCG2* promoter regions were amplified and cloned into the *KpnI /BamHI* sites of the pGreenII 0800-LUC (pLUC) vector to generate PstABCG2:LUC reporter constructs. The ORF of *PstGATA* was amplified and cloned into the *SacI/SpeI* sites of the pGreenII 62-SK (pSK) vector to generate the 35S: PstGATA construct. Both were verified by restriction digestion and sequencing.

The primers used for all constructs are listed in Supplemental Table 2.

### Bioinformatics Analysis of Transcriptome

To investigate the effects of triadimefon and flubeneteram on *Pst*, samples from both treatments (n = 30) were combined as the experimental group and compared with a control group (n = 15). Differentially expressed genes (DEGs) were identified using DESeq2 with the criteria |log₂FC| > 2 and adjusted *P* < 0.05. Principal component analysis (PCA) confirmed no significant differences in global transcriptional patterns between the two fungicide treatments, thereby validating their combination for subsequent analyses. KEGG pathway enrichment analysis was performed, followed by gene set enrichment analysis (GSEA) across all predefined KEGG pathways. The top 20 enriched pathways were visualized.

Candidate ATP-binding cassette (ABC) transporter genes were identified genome-wide using HMMER searches against PF00005 and PF00664 domains, referencing known ABC transporters from *Puccinia graminis* and *Saccharomyces cerevisiae*. Candidate genes were further screened using BLASTP (e-value < 1e-5) in TBtools-II. Phylogenetic analysis was conducted in MEGA 11 with the neighbor-joining algorithm, and the resulting phylogenetic tree was visualized via Chiplot (https://www.chiplot.online/). Venn diagrams comparing DEGs with ABC transporter candidates were generated using the matplotlib_venn library in Python. Time-series transcriptome data were integrated using OmicsTIDE (https://omicstide-tuevis.cs.uni-tuebingen.de/). The optimal cluster number (*k* = 6) for *k*-means clustering was determined based on silhouette scores. Genes showing consistent expression patterns across both fungicide treatments were selected for functional validation. RNA-seq data generated in this study are deposited in the China National GeneBank under accession numbers CNP0007780 and CNP0007841.

### Barley stripe mosaic virus-mediated gene silencing

To assess the role of *PstABCG2* in fungicides resistance, two unique cDNA fragments of *PstABCG2* were selected for BSMV-mediated host-induced gene silencing (HIGS) following established protocols [40]. These fragments showed no homology to wheat genomic sequences (BLAST analysis, URGI database). The wheat plants inoculated with BSMV-TaPDS (phytoene desaturase) were used as the positive control, whereas the BSMV-γ-inoculated plants were acted as the negative control. After viral inoculation, Suwon 11 seedlings were maintained in a growth chamber (25-27 °C, 9-12 d). Upon observing the characteristic photobleaching phenotype in *BSMV-TaPDS*-treated plants, fourth leaves were challenge-inoculated with a low-resistance *Pst* isolate. HIGS assays were conducted as described [372], with three independent biological replicates.

### Histological observation of fungal growth

To evaluate the function of *PstABCG2* under fungicides stress, fungal growth was monitored microscopically. Fungicides solution (100 μg/mL, 97.5% active ingredient, dissolved in acetone) were applied to wheat leaves at 3 days post-inoculation (dpi), with acetone-only treatment serving as the control [6]. Leaf segments (2 cm) were collected at 72 hours post-treatment (hpt) and processed for fluorescence microscopy as described by Liu et al. [41]. Hyphae were stained with WGA-Alexa488 (Invitrogen) and visualized using an Olympus BX53F microscope (450-480 nm excitation/515 nm emission). Fungal colonization was quantified by assessing hyphal growth at 50 randomly selected infection sites per treatment.

### Generation and identification of wheat transgenic plants

The wheat variety Fielder was used to generate transgenic plants via Agrobacterium-mediated transformation. A specific *PstABCG2* cDNA fragment was cloned into the pC336 (Ubi:GWRNAi:Nos) vector using Gateway technology and transformed into *Agrobacterium tumefaciens* strain EHA105, following Aadel et al. [42]. Phenotypic evaluations were performed using protocols consistent with HIGS experiments [43]. Specifically, 15-20 inoculated leaves were examined for phenotypic characterization and urediniospore quantification. Infection areas were measured from 50 infection sites on three leaves, and RNA was extracted from three separate leaves to assess gene silencing efficiency.

### Gene overexpression in *F. graminearum*

The fungal pathogen *Fusarium graminearum* was utilized to generate overexpression strains of *PstABCG2* and its mutant variant *PstABCG2 ^E1184Y^*. For genetic transformation, these genes were introduced into protoplasts of the *F. graminearum* strain 04580, specifically a ΔFgABCG2 knockout mutant lacking the homologous FgABCG2 gene. Fungal protoplast preparation and subsequent polyethylene glycol (PEG)-mediated transformation were carried out following established protocols [44]. Transformants were screened via RT-PCR using gene-specific primers and validated by Western blotting. Positive strains were isolated for further analyses.

### Real-time PCR

To evaluate *PstABCG2* expression under both control and fungicides-treated conditions, samples were collected from wheat cultivar MX169 infected with *Puccinia striiformis* isolate. Urediniospores and infected leaves were harvested at 6, 12, 24, 48, 72 and 120 post-inoculation (hpi). Additionally, for transcript level analysis, leaves of wheat cultivar Su11 inoculated with *Pst* were sampled at 48, 72, and 120 hpi. Quantitative reverse transcription PCR (qRT-PCR) was conducted using a CFX Connect Real-Time System (Bio-Rad, Hercules, CA, USA). Each 25 μL reaction mixture contained: 12.5 μL of LightCycler® SYBR Green I Master Mix, 2 μL of diluted cDNA (1:5), 8.9 μL of nuclease-free H₂O and 0.8 μL each of forward and reverse primers (10 mM) (see Supplementary Table S2 for primer details). Relative gene expression was calculated using the 2^- ΔΔCT^ method, with normalization to the reference genes *PstEF1* (fungal) and *TaEF-1α*(wheat). Three biological replicates were analyzed per sample, with each PCR run performed in triplicate (three technical replicates).

### RNA-seq data processing and analysis

To examine transcriptional changes in *Pst* in response to triadimefon and flubeneteram treatment, transcriptomic data were collected from infected wheat leaves, with 30 biological replicates per trea tment group. RNA sequencing was conducted by Guangzhou GENE DENOVO Technologies usin g the Illumina HiSeq 4000 platform to generate 150 bp paired-end reads. Raw reads were quality-c hecked using FastQC (http://www.bioinformatics.babraham.ac.uk/projects/fastqc/) and low-quality sequences and adapters were removed with Trimmomatic v0.39 (http://www.usadellab.org/cms/?p age=trimmomatic).

Bioinformatics analysis involved building a reference genome index and aligning quality-filtered reads using HISAT2 v2.4 (http://daehwankimlab.github.io/hisat2/) with default parameters, followed by transcript assembly with StringTie v1.3.1 in a reference-guided manner. Gene expression levels were quantified as FPKM (fragments per kilobase of transcript per million mapped reads) using RSEM, while principal component analysis (PCA) was performed using R statistical packages. Differential expression analysis identified significant genes (*P*<0.05, fold-change>2.0) through DESeq2 v3.7.1(https://bioconductor.org/packages/release/bioc/html/DESeq2.html), with subsequent functional annotation of these DEGs conducted via clusterProfiler for KEGG pathway and Gene Ontology (GO) enrichment analyses.

### Protein expression in *E. coli* and purification

Recombinant plasmid constructs were transformed into *E. coli* BL21 competent cells for protein expression. Following transformation, bacterial cultures were induced with 0.5 mM IPTG and grown for 14 hours at 16°C with constant shaking at 250 rpm. Harvested cells were subsequently disrupted via sonication in lysis buffer (50 mM Tris-HCl, 150 mM NaCl, pH 7.5). The crude lysate was then subjected to affinity purification using Ni-NTA beads (GE Healthcare, #45-002-986), with target proteins eluted using a buffer containing 200 mM imidazole. The eluted protein fractions were dialyzed at 4°C for 2 hours to remove imidazole, followed by centrifugation at 4°C to eliminate particulate contaminants. Finally, the purified protein was concentrated using size-exclusion chromatography and analyzed by SDS-PAGE to verify purity and molecular weight.

### In vitro iron affinity assay

The binding affinities between PstABCG2 and two fungicides were analyzed using microscale thermophoresis (MST) and isothermal titration calorimetry (ITC), following established methodologies [45–46]. For MST analysis, target proteins at 100 nM concentration were fluorescently labeled using the Monolith His-tag labeling kit RED-tris-NAT 2nd generation (NanoTemper Technologies) through a 30-minute incubation at room temperature. Serial dilutions of triadimefon (initial concentration: 0.1 μg/mL) were prepared in 16 PCR tubes using a 10-fold dilution gradient, with 10 μL of labeled protein added to each tube. Measurements were conducted on a Monolith NT instrument (NanoTemper Technologies GmbH) following manufacturer’s protocols, with binding affinities calculated using MO Affinity Analysis software (v2.3). For ITC measurements, a 200 μL syringe was loaded with 0.1 μg/mL triadimefon and flubeneteram solution, while the sample cell contained 500 nM protein solution. Titrations were performed with 5 μL injections at 300-second intervals, totaling 50 injections under constant stirring. Control experiments determining the heat of dilution for both protein and water were conducted to enable proper baseline correction of binding isotherms. All ITC data were processed and analyzed using Nano Analyze software.

### Yeast One-Hybrid (Y1H) assay

The yeast 1-hybrid assay was performed following as described [47]. The bait plasmid was linearized and integrated into the Y1HGold yeast genome via homologous recombination, followed by selection on SD/-Ura plates. Successful integrants were confirmed by PCR and tested for AbA sensitivity to determine minimal inhibitory concentration (typically 200-400 ng/mL). The prey plasmid was transformed into yeast containing the integrated bait using the lithium acetate/PEG method, with transformants selected on SD/-Leu plates. Positive colonies were screened on SD/-Leu plates containing the predetermined AbA concentration to detect potential interactions. All cloning steps utilized *E. coli* DH5α for plasmid propagation, with standard molecular biology techniques for purification and verification.

### Firefly luciferase reporter assay

The firefly luciferase reporter assay was conducted according to established protocols [48]. Briefly, recombinant reporter constructs were transformed into *Agrobacterium tumefaciens* strain GV3101 for plant infiltration. Forty-eight hours after agroinfiltration, the transfected leaves were treated with 0.1 mM luciferin solution and dark-adapted for 5 minutes to allow substrate penetration. Luminescence signals were detected using a high-sensitivity CCD camera system (CHEMIPROHT 1300B/LND, 16-bit; Roper Scientific) under low-light conditions. Acquired luminescence images were subsequently analyzed and quantified using ImageJ software (NIH) for comparative expression studies.

### Electrophoretic mobility shift assay

The Electrophoretic mobility shift assay was conducted according to established protocols [49] Briefly, potential PstGATA transcription factor-binding cis-elements in the PstABCG2 promoter were predicted using the Jasper database (probe sequences provided in Supplemental Table S2). Three biotin-labeled probes (P1, P2, P3) containing predicted *cis*-regulatory motifs were synthesized by Shanghai Sangon Biotech and subsequently labeled using the EMSA Probe Biotin Labelling Kit (Beyotime). Unlabeled oligonucleotides served as competitors. Binding reactions were performed using the LightShift Chemiluminescent EMSA Kit (Thermo Scientific, USA) following manufacturer’s protocols, with all experiments conducted in three biological replicates.

### Molecular docking analysis

Tertiary structures of PstABCG2, PstABCG2^E1184Y^, and FgABCG2 (*Fusarium graminearum*) were predicted using AlphaFold3. Docking with triadimefon was simulated in AutoDock Vina to identify binding residues and docking scores. For FgABCG2, site-directed saturation mutagenesis at Vina-predicted residues was performed using FoldX to assess effects on protein stability and ligand-binding affinity. Docking results and protein-ligand interactions were visualized in PyMOL.

Key residues for binding triadimefon and flubeneteram to PstABCG2 and PstABCG2^E1184Y^ were predicted with AlphaFold3 (https://alphafoldserver.com). Preliminary docking was performed using CB-Dock2 (https://cadd.labshare.cn/cb-dock2/php/index.php). Saturation mutagenesis of residues N1132, E1184, and W885 was conducted using FoldX PositionScan to calculate ΔΔG changes upon mutation to all 20 amino acids. Final docking simulations in CB-Dock2 compared binding free energy (ΔG), intermolecular interactions (hydrogen bonds, halogen bonds, π–π stacking, anion–π interactions), and interaction distances (Å).

### Statistical analyses

Statistical analyses were performed using DPS software (version 7.05), with Duncan’s multiple range test applied to determine significant differences among treatments. For triadimefon sensitivity assessment, EC50 values were log-transformed and subjected to Spearman’s rank correlation analysis. All data are presented as means ± standard deviation. Significant differences (*P* < 0.01) were indicated in figures using asterisks, with the following significance thresholds: \**P* < 0.01, \*\**P*< 0.001, and \*\*\**P* < 0.0001.

## Supporting information

**S1 Table. Viral protein sequences used for phylogenetic reconstruction.**

**S2 Table. Primers used in this study.**

**S1 Fig. PCA and differential expression analysis of *Pst* transcriptome under fungicide treatments. (A)** PCA plot showing flueneteram-treated samples (95% confidence ellipse) completely encompassed within the ellipse of triadimefon-treated samples. **(B) V**olcano plot displaying differentially expressed genes (DEGs) between combined fungicide-treated groups (triadimefon + flueneteram, n=30) and control group (n=15), with significance thresholds set at |log2FC| > 2 and *P* < 0.05.

**S2 Fig. Expression analysis of four ABC transporter G family genes (PstABCG2, PstABCG6, PstABCG7 and PstABCG8) in *Pst* under control and fungicide treatment conditions.** Expression analysis of four ABC transporter G family genes in *Pst* at 24 **(A)** and 48 hours **(B)** after treatment with triadimefon and fluindapyr. Wheat seedlings (cv. Mingxian 169) were inoculated with *Pst* urediniospores at 10 days after planting and treated with triadimefon and flueneteram at 3 days after *Pst* inoculation. Staring 3 days after treated with triadimefon and flueneteram, wheat leaves were collected at 0, 24, 48, 72, and 120 h post triadimefon treatment for RNA-sequencing to determine the expression levels of the candidate genes. Expression levels were normalized based on the *Pst* elongation factor gene (*PstEF*). The relative expression values of four ABC transporter G family genes were calculated using the comparative threshold method (2^−ΔΔCt^). \*\*\*\**p*<0.001, \*\*\**p*<0.001, \*\**p*<0.01 and \**p*<0.1.

**S3 Fig. Silence of *PstABCG2* enhances the sensitivity of *Pst* to triadimefon and flueneteram by BSMV-HIGS. (A)** Disease phenotype of *Pst* inoculated with the fourth leaves of Sumon 11 that fully displayed viral symptoms under both triadimefon and flueneteram treatments. **(B)** Silencing efficiency assay of *PstABCG2* in the HIGS plants under triadimefon and flueneteram treatments. RNA samples were isolated from the fourth leaves of knock-down and control plants at 24, 48 and 72 hpi of triadimefon and flueneteram treatments. The *PstEF* gene was used as an internal control. **(C)** Urediniospore biomass on wheat leaves treated with triadimefon and flueneteram at 10 days post-treatment. **(D)** Fluorescence microscopy observation of mycelial growth of *Pst* in wheat leaves after silencing of *PstABCG2* after 72 hpi of triadimefon and flueneteram treatment. **(E)** Comparison of the mycelial area of each infected unit in wheat leaves treated with triadimefon and flueneteram at 72 hpi. Values represent the mean ± standard deviation of three independent samples with 60 infection sites. Single asterisk (*P* < 0.05) and double asterisks (*P* < 0.01) indicate a significant difference from the water control based on the Student’s *t*-test.

**S4 Fig. Purification and immunoblot verification of recombinant PstABCG2 and PstABCG2^E1184Y^ proteins. (A)** SDS-PAGE analysis of PstABCG2 purification (Lane M: Protein marker; Lane 1: PstABCG2). **(B)** SDS-PAGE analysis of PstABCG2 ^E1184Y^ purification (Lane M: Protein marker; Lane 1: PstABCG2 ^E1184Y^). **(C)** Western blot of purified PstABCG2 (Anti-His tag). **(D)** Western blot of purified PstABCG2 ^E1184Y^ (Anti-His tag).

## Author Contributions

**Conceptualization:** Fan Ji

**Data curation:** Fan Ji, Xin-Pei Gao, Ya-Ning Liu

**Formal analysis:** Fan Ji, Xin-Pei Gao, Ya-Ning Liu

**Investigation:** Fan Ji, Xin-Pei Gao, Ya-Ning Liu, Ying Li

**Methodology:** Fan Ji, Xin-Pei Gao, Ya-Ning Liu, Ying Li **Validation:** Fan Ji, Ying Li

**Visualization:** Fan Ji, Ya-Ning Liu

**Writing – original draft:** Fan Ji

**Writing – review & editing:** Li-Li Huang, Gang-Ming Zhan, Zhen-Sheng Kang

## References

1. Tang TF, Tian SL, Wang HY, Lv XT, Xie YL, Liu JY, et al. Wheat-RegNet: An encyclopedia of common w heat hierarchical regulatory networks. Mol Plant. 2023; 16(2):318–321. 10.1016/j.molp.2022.12.018 PMID: 36575798

2. Chen WQ, Colin W, Chen XM, Kang ZS, Liu TG. Wheat stripe (yellow) rust caused by *Puccinia striiformis* f. sp. *tritici*. Mol Plant Pathol. 2014; 15(5):433–446. 10.1111/mpp.12116 PMID: 24720631

3. Chen XM. Pathogens which threaten food security: *Puccinia striiformis*, the wheat stripe rust pathogen. Foo d Security. 2020; 12(2):239–251. 10.1007/s12571-020-01016-z PMID: 32214664

4. Zhao J, Kang ZS. Fighting wheat rusts in China: a look back and into the future. Phytopathol Res. 2023; 5(2): 6–7. 10.1186/s42483-023-00159-z PMID: 36865360

5. Kang ZH, Li X, Wan AM, Wang MN, Chen XM. Differential sensitivity among *Puccinia striiformis* f. sp. *tritici* isolates to propiconazole and pyraclostrobin fungicides. Can J Plant Pathol. 2019; 41(3):415–434. 10.1080/07060661.2019.1577301 PMID: 30915827

6. Zhan GM, Ji F, Zhao J, Liu Y, Zhou AH, Xia MH, et al. Sensitivity and resistance risk assessment of *Puccinia striiformis* f. sp. *tritici* to triadimefon in China, Plant Dis. 2022; 106(6):1690–1699. 10.109 4/PDIS-10-21-2168-RE PMID: 34962420

7. Wu XX, Bian Q, Lin QJ, Sun Q, Ni XY, Xu XF, et al. Sensitivity of *Puccinia striiformis* f. sp. *tritici* isolates from China to triadimefon and cross-resistance against diverse fungicides. Plant Dis. 2020; 104(8):2082–2085. 10.1094/PDIS-01-20-0009-RE PMID: 3255228

8. Ji F, Zhang JT, Chen XM, Liu BF, Zhou AH, Feng YX, et al. Effects of flubeneteram on inhibiting the devel opment of *Puccinia striiformis* f. sp. *tritici* in wheat leaves. J Agric Food Chem. 2023; 71(13):5162–5171. htt ps://doi.org/10.1021/acs.jafc.3c00499 PMID: 36946748

9. Maxfield FR, Tabas I. Role of cholesterol and lipid organization in disease. Nature. 2005; 438(7068):612–621. 10.1038/nature04399 PMID: 16319881

10. Geißel B, Loiko V, Klugherz I, Zhu Z, Wagener N, Kurzai O, et al. Azole-induced cell wall carbohydrate patches kill *Aspergillus fumigatus*. Nat Commun. 2018; 9(1):3098. 10.1038/s41467-018-05497-7 PMID: 30082817

11. Chen SN, Yuan NN, Schnable G, Luo CX. Function of the genetic element ’Mona’ associated with fungicide resistance in *Monilinia fructicola*. Mol Plant Pathol. 2016; 18(1):90–97. 10.1111/mpp.12387 P MID: 26918759

12. Liu ZY, Jian YQ, Chen Y, Kistler HC, He P, Ma ZH, et al. A phosphorylated transcription factor regulates st erol biosynthesis in *Fusarium graminearum*. Nat Commun. 2019; 10(1):1228. 10.1038/s4146 7-019-09145-6 PMID: 30874562

13. Tucker MA, Lopez RF, Mullins JGL, Jayasena K, Oliver RP. Analysis of mutations inWest Australian popul ations of *Blumeria graminis* f. sp. *hordei* Cyp51 conferring resistance to DMI fungicides. Pest Manag Sci. 2020; 76(4):1265–1272. 10.1002/ps.5636 PMID: 31595590

14. Zeng LQ, Luo RT, Chen Q, Hao GF, Zhu XL, Yang GF, et al. Discovery of fungicide flubeneteram. J Pestic Sci. 2022; 24(5):895–903. 10.16801/j.issn.1008-7303.2022.0028

15. Chen ZL, Shi TL, Zhang L, Zhu PL, Deng MY, Huang C, et al. Mammalian drug efflux transporters of the ATP binding cassette (ABC) family in multidrug resistance: a review of the past decade. Cancer Lett. 2016; 370(1):153–164. 10.1016/j.canlet.2015.10.010 PMID: 26499806

16. Jenness MK, Carraro N, Pritchard C, Murphy AS. The *Arabi dopsis* ATP-binding cassette transporter ABCB 21 regulates auxin levels in cotyledons, the root pericycle, and leaves. Front Plant Sci. 2019; 10(1):806. 10.3389/fpls.2019.00806 PMID: 31275345

17. Nijland M, Lefebvre SN, Thangar C, Slotboom DJ. Bidirectional ATP-driven transport of cobalamin by the mycobacterial ABC transporter BacA. Nat Commun. 2024; 9(2):17. 10.1038/s41467-024-469 17-1 PMID: 38521790

18. Firoz MJ, Xiao X, Zhu FX, Fu YP, Jiang DH, Schnabelc G, et al. Exploring mechanisms of resistance to di methachlone in *Sclerotinia sclerotiorum*. Pest Manag Sci. 2016; 72(4):770–779. 10.1002/ps.40 51 PMID: 26037646

19. Mosbach A, Leroch M, Kretschmer M, Mernke D, Walker AS, Fill IS, et al. *Botrytis cinerea* mutations in trans cription factor *Mrr1* leading to overexpression of efflux transporter AtrB and multiple drug resistance in *Bo trytis cinerea* field strains. Julius-Kühn-Archiv. 2010; 428(4):226.

20. Alekshun MN, Levy SB. Molecular mechanisms of antibacterial multidrug resistance. Cell. 128(6):1037–50. 10.1016/j.cell.2007.03.004 PMID: 17382878

21. Gupta AK, Wang T, Mann A, Piguet V, Chowdhary A, Bakotic WL. Mechanisms of resistance against allyl amine and azole antifungals in Trichophyton: A renewed call for innovative molecular diagnostics in suscept ibility testing. PLoS Pathog. 2025; 21(2):e1012913. 10.1371/Journal.ppat.1012913 PMID: 39 932950

22. Hu P, Liu YC, Zhu XL, Kang HX. ABCC Transporter gene *MoABC-R1* is associated with pyraclostrobin tol erance in *Magnaporthe oryzae*. J Fungi (Basel). 2023; 9(9):917. 10.3390/jof9090917 PMID: 37755025

23. Cheng XK, Zhang JT, Liang ZY, Wu ZC, Liu PF, Hao JJ, et al. Multidrug resistance of Rhizoctonia solani d etermined by enhanced efflux for fungicides. Pestic Biochem Physiol. 2023; 195:105525. 10.1016/j.pestbp.2023.105525 PMID: 37666584

24. Wang J, Xiao CW, Liang S, Noman M, Cai YY, Zhang Z, et al. Comparative functional analysis of a new C DR1-like ABC transporter gene in multidrug resistance and virulence between *Magnaporthe oryzae* and *Tric hophyton mentagrophytes*. Cell Commun Signal. 2025; 23(1):69. 10.1186/s12964-024-02022-w PMID: 39920659

25. Bardas GA, Veloukas T, Koutita O, Karaoglanidis GS. Multiple resistance of *Botrytis cinerea* from kiwifruit to SDHIs, QoIs and fungicides of other chemical groups. Pest Manag Sci. 2010; 66(9):967–973. https://doi.o rg/10.1002/ps.1968 PMID: 20730988

26. Cheng XK, Man XJ, Wang ZT, Liang L, Zhang F, Wang ZW, et al. 2020. Fungicide SYP-14288 inducing m ultidrug resistance in *Rhizoctonia solani*. Plant Dis. 2020; 104(10):2563–2570. 10.1094/PDIS-01-20-0048-RE PMID: 32762501

27. Leroux P, Walker AS. Activity of fungicides and modulators of membrane drug transporters in field strains of *Botrytis cinerea* displaying multidrug resistance. Eur J Plant Pathol 2013; 135:683–693. 10.1007/s10658-012-0105-3

28. Moye-Rowley WS. Transcriptional control of multidrug resistance in the yeast Saccharomyces. Prog Nucleic Acids Res Mol Biol. 2003; 73:251–279. 10.1016/s0079-6603(03)01008-0 PMID: 12882520

29. Yamano S, Tsukuda Y, Mizuhara N, Yamaguchi Y, Ogita A, Fujita KI. Dehydrozingerone enhances the fun gicidal activity of glabridin against *Saccharomyces cerevisiae* and *Candida albicans*. Lett Appl Microbiol. 2 023;76(4):ovad040. 10.1093/lambio/ovad040 PMID: 36990694

30. Li YQ, Dai MY, Lu L, Zhang YW. The C2H2-Type transcription factor ZfpA, coordinately with CrzA, affe cts azole susceptibility by regulating the multidrug transporter gene atrF in *Aspergillus fumigatus*. Microbiol Spectr. 2023; 11(4):e00325–23. 10.1128/spectrum.00325-23 PMID: 37318356

31. Kretschmer M, Leroch M, Mosbach A, Walker AS, Fillinger S, Mernke D, et al. Fungicide-driven evolution and molecular basis of multidrug resistance in field populations of the Grey mould fungus *Botrytis cinerea*. PLoS Pathogens. 2009; 5(12):e1000696. 10.1371/Journal.ppat.1000696 PMID: 20019793

32. Sang H, Hulvey JP, Green R, Xu H, Im J, Chang TY, et al. A xenobiotic detoxification pathway through tran scriptional regulation in filamentous fungi. MBio. 2018; 9(4):e418–e457.10.1128/mBio.00457-18 PMID: 30018104

33. Wu ZC, Zhang JT, Zhang BR, Yu CX, Wang ZW, Hao ZG, et al. Regulatory role of the transcription factor BcMr1 in efflux-mediated multidrug resistance in *Botrytis cinerea* via pre-adaptation to plant secondary met abolites. Int J Biol Macromol. 2025; 311(1):143520. 10.1016/j.ijbiomac.2025.143520 PMID: 40306507

34. Pérez-de Los Santos FJ, García-Ortega LF, Robledo-Márquez K, Guzmán-Moreno J, Riego-Ruiz L. Transcri ptome analysis unveils gln3 role in amino acids assimilation and fluconazole resistance in *Candida glabrata*. J Microbiol Biotechnol. 2021; 31(5):659–666. 10.4014/jmb.2012.12034 PMID: 33879640

35. Tian Y, Zhan GM, Chen XM, Angkana T, Lu X, ZhaoJ, et al. Virulence and simple sequence repeat marker segregation in a *Puccinia striiformis* f. sp. *tritici* population produced by selfing a Chinese isolate on *Berberi s shensiana*. Phytopathology. 2016; 106(2):185–191. 10.1094/PHYTO-07-15-0162-R PMID: 26551448

36. Healey A, Furtado A, Cooper T, Henry RJ. Protocol: a simple method for extracting next-generation sequen cing quality genomic DNA from recalcitrant plant species. Plant Methods. 2014; 10:21. 10.1186/1746-4811-10-21 PMID: 25053969

37. Liu C, Wang YQ, Wang YF, Du YY, Song C, Song P, et al. Glycine-serine-rich effector *PstGSRE4* in *Pucci nia striiformis* f. sp. *tritici* inhibits the activity of copper zinc superoxide dismutase to modulate immunity in wheat. PLoS Pathog. 2022; 18(7):e1010702. 10.1371/Journal.ppat.1010702 PMID: 35881621

38. Duan WL, Hao ZK, Pang HH, Peng YX, Xu YW, Zhang YF, et al. Novel stripe rust effector boosts the trans cription of a host susceptibility factor through affecting histone modification to promote infection in wheat. New Phytol. 2024; 241(1):378–393. 10.1111/nph.19312 PMID: 37828684

39. Xu HJ, Ye M, Xia AL, Jiang H, Huang PP, Liu HQ, et al. The Fng3 ING protein regulates H3 acetylation an d H4 deacetylation by interacting with two distinct histone-modifying complexes. New Phytol. 2022; 235(6): 2350–2364. 10.1111/nph.18294 PMID: 35653584

40. Holzberg S, Brosio P, Gross C, Pogue GP. Barley stripe mosaic virus-induced gene silencing in a monocot plant. Plant J. 2002; 30(3):315–327 (2002). 10.1046/j.1365-313x.2002.01291.x PMID: 12000679

41. Liu J, Guan T, Zheng PJ, Chen LY, Yang Y, Huai BY, et al. An extracellular Zn-only superoxide dismutase from *Puccinia striiformis* confers enhanced resistance to host-derived oxidative stress. Environ. Microbiol. 2 016; 18(11):4118-4135. 10.1111/1462-2920.13451. PMID: 27399209

42. Aadel H, Abdelwahd R, Udupa SM, Diria G, El Mouhtadi A, Ahansal K, et al. Agrobacterium-mediated transformation of mature embryo tissues of bread wheat (*Triticum aestivum* L.) genotypes. Cereal Res Commun. 2018; 46(1):10–20. 10.1556/0806.45.2017.055

43. Xu Q, Tang CL, Wang XD, Sun ST, Zhao JR, Kang ZS, et al. An effector protein of the wheat stripe rust fun gus targets chloroplasts and suppresses chloroplast function. Nat Commun. 2019; 10(1):5571. 10.1038/s41467-019-13487-6 PMID: 31804478

44. Jiang C, Hei RN, Yang Y, Zhang SJ, Wang QH, Wang W, et al. An orphan protein of *Fusarium graminearum* modulates host immunity by mediating proteasomal degradation of TaSnRK1α. Nat Commun. 2020; 11(1) :4382. 10.1038/s41467-020-18240-y PMID: 32873802

45. Zhang JF, Li CX, Wang GD, Cao JW, Yang X, Liu XB, et al. α-Amylase inhibition of a certain dietary poly phenol is predominantly affected by the concentration of α-1, 4-glucosidic bonds in starchy and artificial sub strates. Food Res Int. 2022; 157:111210. 10.1016/j.foodres.2022.111210 PMID: 35761532

46. Yu H, Zhao Q. Profiling additional effects of aptamer fluorophore modification on microscale thermophores is characterization of aptamer–target binding. Anal Chem. 2023; 95(46):17011–17019. 10.1021/acs.analchem.3c03603 PMID: 37946406

47. Liu D, Li YY, Zhou ZC, Xiang X, Liu X, Wang J, et al. Tobacco transcription factor bHLH123 improves sal t tolerance by activating NADPH oxidase NtRbohE expression. Plant Physiol. 2021; 186(3):1706–1720. 10.1093/plphys/kiab176 PMID: 33871656

48. Du YW, Jia HC, Yang Z, Wang SH, Liu YY, Ma HY, et al. Sufficient coumarin accumulation improves apple resistance to Cytospora mali under high-potassium status. Plant Physiol. 2023; 192(2):1396–1419. 10.1093/plphys/kiad184 PMID: 36943289

49. Du YW, Liu GL, Jia HC, Liu Y, Tan Y, Wang SH, et al. Changes in planta K nutrient content altered the interaction pattern between *Nicotiana benthamiana* and *Alternaria longipes*. Plant Cell Environ. 2024; 47(9):3619–3637. 10.1111/pce.14956 PMID: 38747645

